# The role of prolines and glycine in the transmembrane domain of LAT

**DOI:** 10.1101/2020.08.10.244251

**Authors:** Daniela Glatzová, Harsha Mavila, Maria Chiara Saija, Tomáš Chum, Lukasz Cwiklik, Tomáš Brdička, Marek Cebecauer

## Abstract

LAT is a critical regulator of T cell development and function. It organises signalling events at the plasma membrane. However, the mechanism, which controls LAT localisation at the plasma membrane is not fully understood. Here, we studied the impact of helix-breaking amino acids, two prolines and one glycine, in the transmembrane segment on localisation and function of LAT. Using *in silico* analysis, confocal and superresolution imaging and flow cytometry we demonstrate that central proline residue destabilises transmembrane helix by inducing a kink. The helical structure and dynamics is further regulated by glycine and another proline residue in the luminal part of LAT transmembrane domain. Replacement of these residues with aliphatic amino acids reduces LAT dependence on palmitoylation for sorting to the plasma membrane. However, surface expression of these mutants is not sufficient to recover function of non-palmitoylated LAT in stimulated T cells. These data indicate that geometry and dynamics of LAT transmembrane segment regulate its localisation and function in immune cells.

## INTRODUCTION

Function of proteins (e.g., enzymes or signal transducers) is defined by their structure, but also localisation within a cell or an organism. Amino acid sequence, small molecule co-factors and interacting partners are primary determinants of the structure. However, features defining protein localisation are less well understood. The process defining protein localisation is called sorting. Short motifs within a primary sequence were found to firmly control sorting of proteins to some organelles, e.g., the ER retention (KDEL motif), the ER exit (DxE and YxxΦ coupled motifs) or transport to/from the nucleus (NES/NLS motifs) [1–3]. Trafficking of proteins to the plasma membrane appears to be controlled by several unrelated determinants. Post-translational modifications, such as glycosylation and palmitoylation facilitate anterograde membrane transport of proteins to the cell surface [4–6]. In the opposite direction, ubiquitination regulates retrograde protein transport from the surface towards the cell interior [7]. However, the primary determinant of the plasma membrane sorting, at least for single-spanning membrane proteins, is the length and amino acid composition of the transmembrane domain (TMD) [6, 8]. A comprehensive study demonstrated that, in vertebrates, proteins with TMDs of 22 residues or longer localise to the plasma membrane [9]. Those with shorter TMDs remain in the Golgi apparatus or the ER. The voluminous amino acids (Phe, Tyr, Trp) were found to accumulate in the cytosolic part of the TMD of the plasma membrane proteins. The position of other amino acids is also non-random [9, 10]. These data indicate that physico-chemical properties of TMDs strongly influence trafficking of proteins in cells.

In vertebrates, the structure of TMD is almost exclusively formed by an α-helix. The hydrophobic core is largely composed of aliphatic (Leu, Val, Ala, Ile) and aromatic amino acids (Phe, Tyr, Trp). Weakly polar amino acids (Ser, Thr, Met, Cys) are also common components of TMDs [10]. Cysteine residues due to their propensity to form disulphide bridges in the oxidative environment, can force the protein to dimerise. Their presence is thus reduced in the exoplasmic part of the TMDs. Moreover, cysteine residues at the cytosolic end of the TMD often represent palmitoylation sites [11]. Positively charged amino acids (Lys, Arg, His) often define the cytosolic end of TMDs (*positive-inside rule*; ref. [12]). In addition, these residues can form electrostatic bridges between the TMDs of multi-spanning or multi-component membrane assemblies [13]. Interestingly, negatively charged amino acids (Asp, Glu) and their amido-derivatives (Asn, Gln) are almost absent from TMDs of vertebrate membrane proteins [10]. However, an increased representation of Asp and Glu has been observed in the membrane-adjacent sequence at the luminal side of the TMD [9, 10]. Finally, proline and glycine are the two amino acids with physico-chemical properties, which can affect the stability of TMDs.

Proline and glycine were defined as typical ‘helix-breakers’ in soluble globular proteins. Indeed, they are virtually absent from the helical structures of such proteins [14]. However, proline is often present in the TMDs of integral membrane proteins, exhibiting the highest occupancy towards the ends but, with a lower frequency, also in the central parts of the α-helices [10, 15, 16]. The cyclic structure of proline makes it unique among the 20 natural amino acids because its amide group lacks the proton necessary for the hydrogen bond stabilising the α-helix or the β-sheet structure [17]. Therefore, prolines in transmembrane (TM) helices are predicted to induce regions of helix distortions and/or dynamic flexibility (i.e., kinks, hinges, swivels; refs. [15, 18]). Such structural distortions may play a role in the transmission of conformational changes along the helix, establish crucial helix-helix packing interactions or geometries, but can also help the helix to adapt the optimal position [15, 19]. Proline was found in the TM helix of many membrane proteins, especially with a polytopic structure [20]. There, the presence of central proline can lead to the change in the helix orientation [16]. Importantly, prolines in TMDs are critical for the function of several proteins [19, 21, 22].

Glycine is also rather frequent in the TMDs of integral membrane proteins [10]. In analogy to proline, glycine helix-affecting properties depend on the local environment [23, 24]. No strong effect of a single glycine residue on basic structure of transmembrane α-helices was found experimentally [23, 24]. It functions primarily as an interface between individual helices of the polytopic membrane proteins and is involved in protein dimerisation [25, 26]. However, when positioned close to the proline, it enhances local dynamics of the helix and, thus, can affect the function of transmembrane peptides or proteins [27, 28]. The complexity of proline and glycine environment-sensitive properties suggests that it is important to characterise their role in the TMD of individual proteins separately.

LAT, linker for activation in T cells, is 36-kDa protein belonging to the family of transmembrane adaptor proteins (TRAPs). It plays an important role in membrane immuno-receptor signalling pathways of T cells, NK cells, mast cells and platelets [29, 30]. LAT is indispensable for the T-cell function and development [31]. It plays an essential role in T-cell activation by providing a platform for signal transmission initiated by the T-cell receptor (TCR) at the plasma membrane [30, 32]. T cells lacking LAT, i.e. the Jurkat mutants J.CaM2.5, fail to mobilise calcium and promote other downstream effector events upon TCR stimulation (e.g., ERK phosphorylation or IL-2 production). Human LAT is a type III transmembrane protein, i.e. it is missing a signal peptide and, thus, uses its unique TMD for its membrane insertion. Its extracellular part has only 3 amino acids, the TMD is 24 residues long and the cytoplasmic tail (235 amino acids) contains nine conserved tyrosine residues involved in T-cell signalling. The cytoplasmic tail of LAT also possesses conserved juxtamembrane cysteines required for its palmitoylation (residues 26 and 29). Palmitoylation is mainly responsible for sorting of LAT to the plasma membrane [6, 33, 34].

In this study, we have investigated the impact of highly conserved proline and glycine residues of the LAT TMD on its plasma membrane localisation and function. We demonstrate that replacement of prolines and glycine with aliphatic amino acids (alanine or leucine) partially recovers the surface localisation of non-palmitoylatable LAT mutant. However, such effect is not sufficient for restoration of its function. We also tested whether nanoscopic surface organisation of LAT mutants is altered and, thus, leads to their malfunction in T cells.

## RESULTS

### Atypical transmembrane domain of LAT contains highly conserved helix-breakers at positions 8, 12 and 17

We have previously shown that TMD controls sorting of LAT to the plasma membrane [6]. General plasma membrane sorting rules determined for other proteins are not obeyed by LAT: i) 24-residues long TMD should provide sufficient sorting signal but LAT, which lacks cysteines at the positions 26 and 29 and, therefore, cannot be palmitoylated, remains trapped in the Golgi apparatus [6, 9, 33], and ii) there are no voluminous amino acids in the exoplasmic portion of the LAT TMD (Fig. 1A). It was previously reported that prolongation of the TMD leads to the plasma membrane localisation of the proteins, which normally reside in the Golgi apparatus [8]. Nevertheless, prolongation of LAT TMD by 6 amino acids (six leucine residues or PILAML sequence) did not overcome dependence of LAT surface expression on its palmitoylation (Supplementary Fig. S1). We observed the accumulation of LAT with prolonged TMD in lysosomes and other unspecified intracellular vesicles (Supplementary Fig. S1). These data indicate that LAT TMD contains residues, which control its geometry or dynamics with respect to the lipid membranes of the secretory pathway.

**Figure 1.**
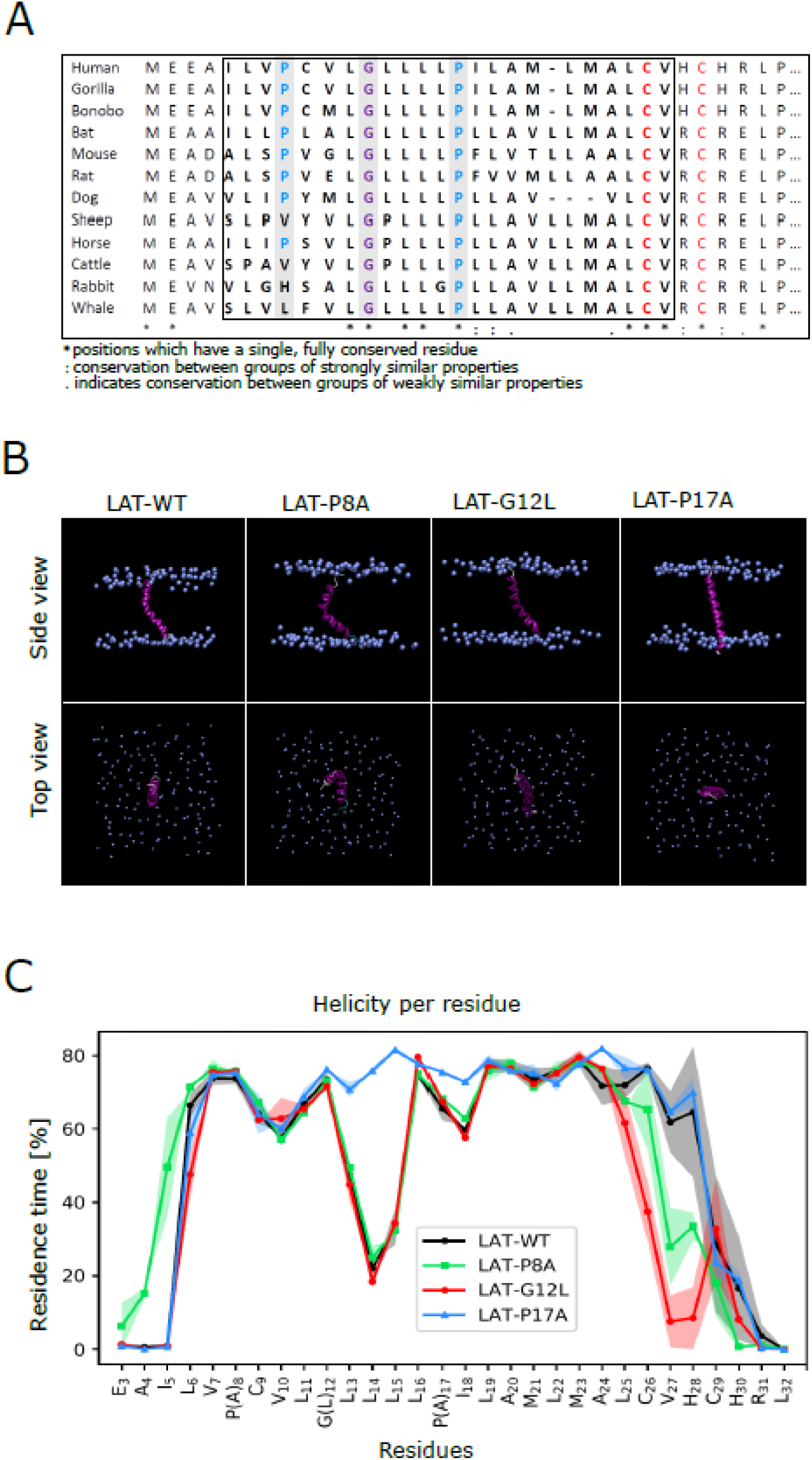
Helix-breaking amino acids in the transmembrane domain of LAT. **(A)** Sequence alignment of LAT transmembrane segment and the adjacent parts (hydrophobic region in bold) showing conservation of amino acids among twelve mammalian species. **(B)** Representative snapshot from MD simulations of LAT TMD peptide and its variants in POPC bilayer. Transmembrane helix is shown in magenta, phospholipid headgroups are indicated with light blue spheres. **(C)** Helicity plot showing the propensity of individual amino acids in the LAT TMD peptide and its variants to accommodate helical geometry. Standard error is indicated as shadowing of the curves.

The amino acid sequence of the LAT TMD exhibits high level of conservation with 48% identity in mammals (Fig. 1A). Among the conserved residues there are two prolines, which were reported to introduce distortions into the helical structure of TMDs in several proteins [15]. One proline is positioned centrally (Pro17) and one close to the luminal end of the LAT TMD (Pro8). Moreover, the luminal part of LAT TMD contains glycine (Gly12) in the position i+4 with respect to Pro8 and i−5 with respect to Pro17. The motif (PxxxG) was described to control the dynamics of pore-forming peptides and proteins [27]. Importantly, helix-breaking residues were shown to exhibit cumulative effect on the structure of TM helices [24]. We, therefore, hypothesised that Pro8, Gly12 and Pro17 residues play a role in membrane stability and sorting of LAT to the plasma membrane.

### All-atom MD simulations indicate kink formation in the helical structure of LAT TMD

We have first characterised the role of Pro8, Gly12 and Pro17 residues in LAT TMD structure using all-atom MD simulations. Residues 2-33, EEAILVPCVLGLLLLPILAMLMALCVHCHRLP, which form the hydrophobic core and membrane proximal parts, were used to simulate human LAT TMD in a lipid membrane composed of 128 palmitoyl-oleoyl phosphatidylcholine (POPC) molecules for 1 μs. The peptide was first enforced to adapt α-helical structure and inserted into the membrane in a transbilayer orientation. Afterwards, the system was left to adjust its optimal conformation for 500 ns. The following 500 ns of the trajectory were used for the analysis of the peptide behaviour in a model membrane. The structure of LAT WT peptide exhibited high level of helicity. However, a kink could be observed in the central part of the peptide at around the position of Leu14 (Fig. 1B,C). This non-helical segment involving residues 13-15 is downstream of Gly12 (i+2 for Leu14) and upstream of Pro17 (i−3 for Leu14). Such deviation within the α-helix is in agreement with previously observed kinks induced by proline and glycine residues in the TMDs of several membrane proteins (e.g., bacteriorhodopsins, ref. [28]). The maximal impact was reported for positions i−3 or i−4 with respect to the proline residue [35]. Maximal effect of glycine was observed for residues i+1 and i+2 [24]. Another deviation from the α-helical structure, even though much smaller compared to the central non-helical segment, could be observed near Pro8 residue (Fig. 1C).

We have further used MD simulations to characterise behaviour of LAT TMD mutants, in which helix-breaking residues were replaced with alanine or leucine. Mutation of proline to alanine in the position 17 (P17A) led to complete disappearance of the kink for the full period of the simulation (Fig. 1B,C). A minor helix deviation near Pro8 was preserved in P17A mutant (Fig. 1C). On the contrary, no effect on the central kink was observed in the mutant P8A. Nevertheless, the mutant exhibited shortening of the helical TM segment (Fig. 1C) and higher dynamics of the kink, as represented by the analysis of the kink angles (Supplementary Fig. S2). The helix was 2 residues shorter compared to LAT WT peptide. No such shortening was observed for P17A mutant (Fig. 1C). MD simulations of G12L mutant of LAT TMD peptide provided similar results as for LAT WT peptide (Fig. 1C and Supplementary Fig. S2). However, the helical TM segment was 3 residues shorter than in the LAT WT peptide. The overall tilt angle of the peptides in the bilayer was similar for all variants except for the G12L mutant, probably caused by shortening of its helical fragment (Supplementary Fig. S3). The *in silico* data thus indicate that Pro17 is critical for the kink formation in the LAT TMD but Pro8 and Gly12 fine-tune the overall TMD structure and dynamics of the kink.

### Mutation of helix-breaking amino acids promotes sorting of non-palmitoylated LAT to the plasma membrane

We have therefore investigated the impact of Pro8, Gly12 and Pro17 on LAT sorting to the plasma membrane. The coding DNA sequence was individually modified to replace Pro8 and Pro17 residues with alanine (Fig. 2A) and expressed as mutant LAT-GFP fusion proteins in LAT-deficient Jurkat T-cell line, J.CaM2.5. Gly12 residue was mutated to leucine to prevent the disruption of leucine-rich segment of the LAT TMD (Figs. 1A and 2A). The cells were transiently transfected with the plasmid DNA and the expression was analysed 16-24h later using confocal scanning microscopy. All mutant proteins exhibited a distribution, which was comparable to the native LAT protein. The proteins prevalently localised to the plasma membrane of transfected J.CaM2.5 cells (Fig. 2B). A partial trapping of the proteins in the Golgi apparatus could also be observed. This observation is in agreement with previous findings that T cells express two pools of LAT, one localized in the plasma membrane and the other in the Golgi apparatus [36]. However, no extensive accumulation of the mutant LAT proteins in intracellular membranes, e.g. the ER, was visible. These results indicate that mutating individual ‘helix-breaking residues’ of the LAT TMD to alanine (or leucine) does not substantially destabilise LAT protein.

**Figure 2.**
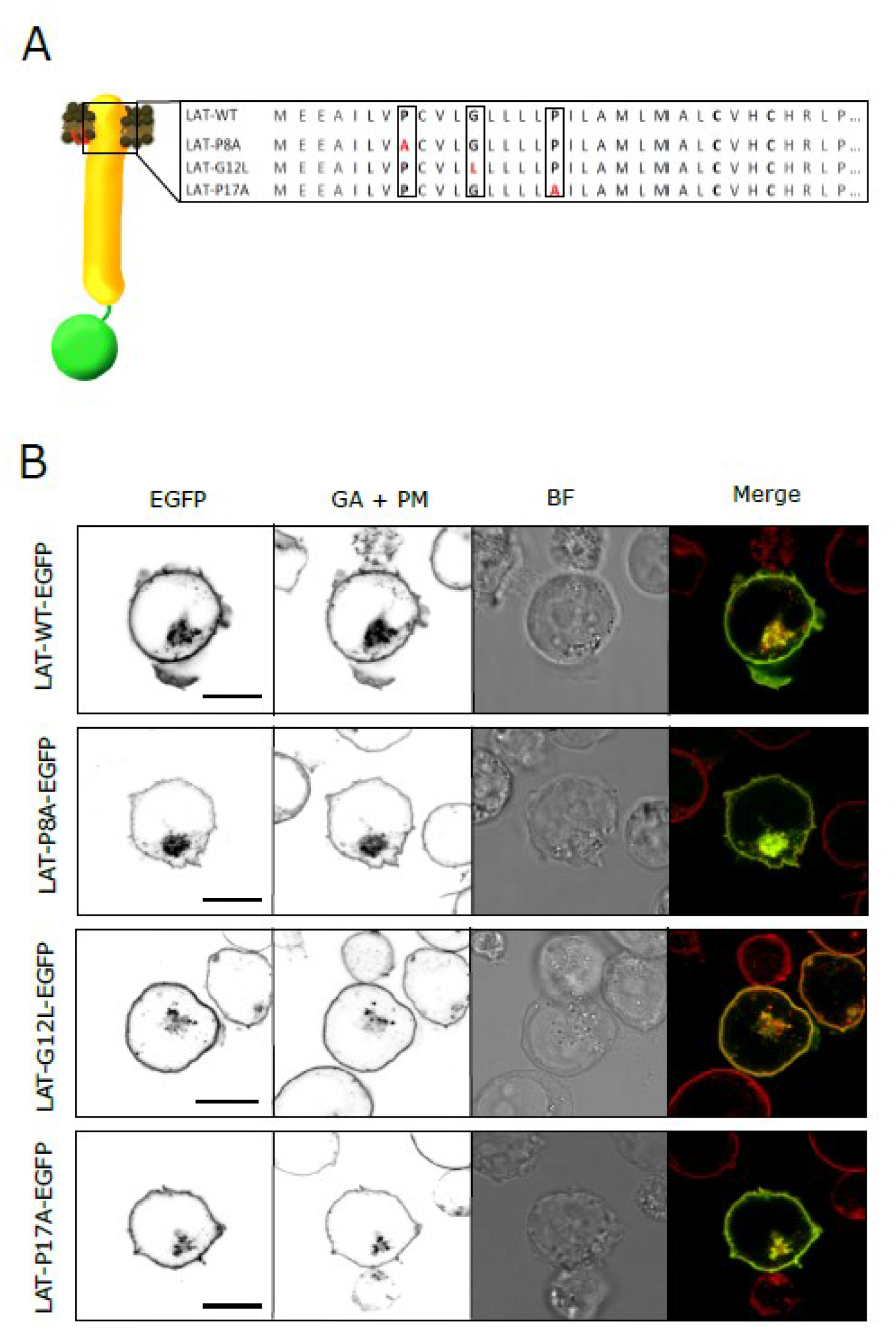
Localisation of palmitoylated LAT variants in J.Cam2.5 T cell line. **(A)** Schematic representation of human LAT construct with attached EGFP and indicated TMD region with accompanied primary sequence. Amino acids mutated in tested LAT variants are highlighted in red. **(B)** Confocal microscopy of J.CaM2.5 T cells expressing LAT-WT-EGFP, LAT-P8A-EGFP, LAT-G12L-EGFP and LAT-P17A-EGFP stained with the plasma membrane (PM) and Golgi apparatus (GA) marker lectin-HPA AF647. Single-channel (EGFP and GA + PM marker) or two-channel overlay images (merge; EGFP – green, GA + PM – red, overlay – yellow) are shown together with a brightfield image (BF). Scale bars, 10 μm. More examples in Supplementary Figs. S4–S7. Representative images of 3 measurement days are shown.

As mentioned above, LAT requires palmitoylation of its membrane proximal cysteines 26 and 29 for its localization to the plasma membrane and function. We were, thus, interested whether a kink in the helix of the TMD determined using MD simulations can influence plasma membrane sorting of LAT in the absence of its palmitoylation. We prepared LAT-GFP variants (Fig. 3A), which combined alanine mutants of Pro8 and Pro17 (or leucine mutant of Gly12) with cysteines 26 and 29 replaced for serines as described before [6]. The mutant proteins were transiently expressed in J.CaM2.5 cells and analysed by confocal microscopy 16-24h post transfection. Strikingly, mutation of helix breaking amino acids in the TMD to alanine or leucine partially restored the plasma membrane localisation of non-palmitoylatable LAT in 30-50% of cells (Fig. 3B and Supplementary Fig. S4–7). Compared to LAT-P8A-CS, slightly stronger surface signal was observed at the plasma membrane in cells expressing LAT-G12L-CS and LAT-P17A-CS (Fig. 3B and Supplementary Figs. S5–7). Nevertheless, LAT-P8A-CS, LAT-P17A-CS and LAT-G12L-CS were also found in the intracellular membranes, mainly Golgi apparatus and the ER. Such intracellular trapping is similar to LAT-CS mutant with preserved kink-forming amino acids (Fig. 3B and Supplementary Fig. S4).

**Figure 3.**
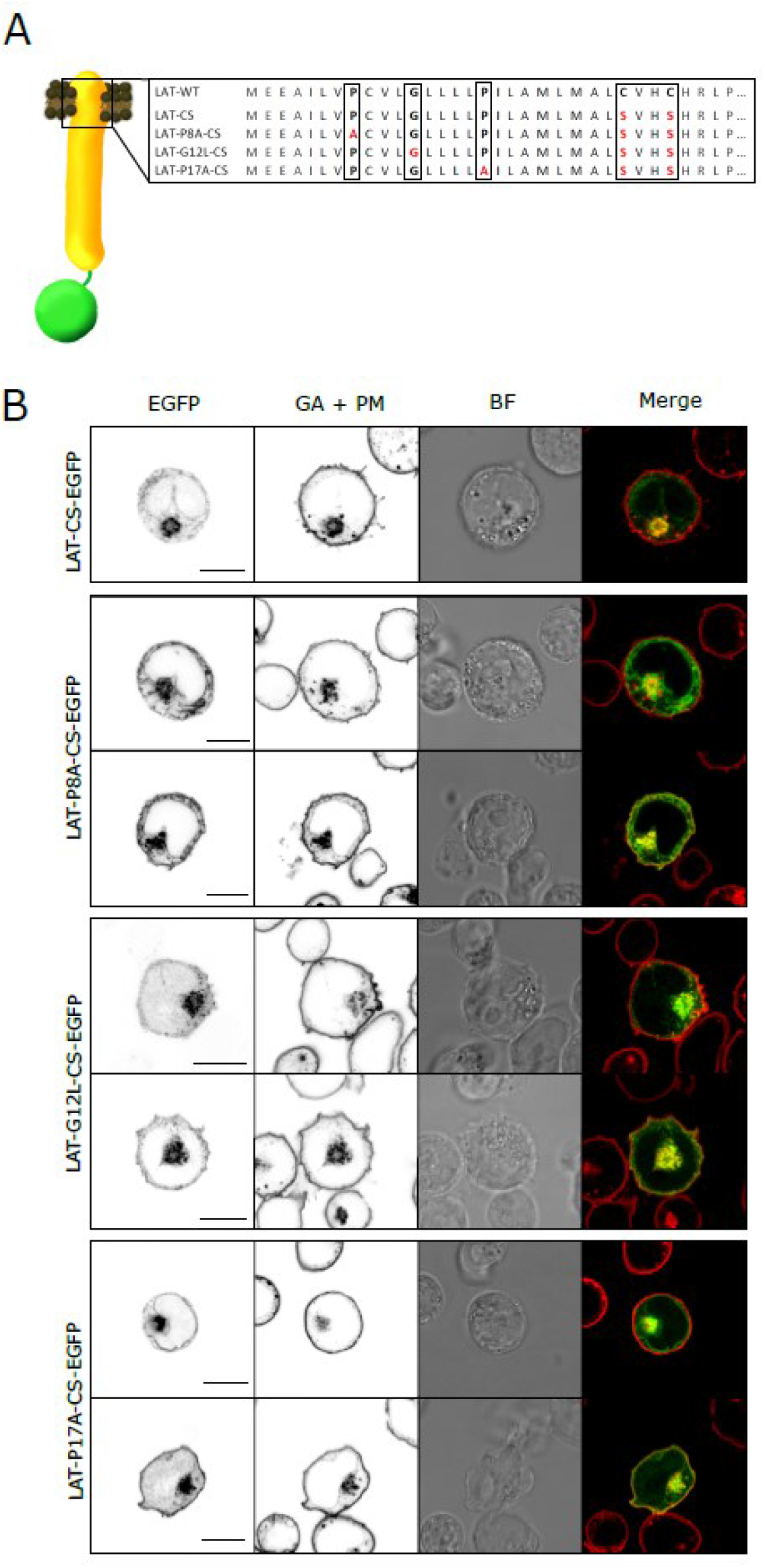
Localisation of non-palmitoylated LAT variants in J.CaM2.5 T cell line. **(A)** Schematic representation of human non-palmitoylated LAT construct with attached EGFP and indicated TMD region with accompanied primary sequence. Amino acids mutated in tested LAT-CS variants are highlighted in red. **(B)** Confocal microscopy of J.CaM2.5 T cells expressing LAT-CS-EGFP, LAT-P8A-CS-EGFP, LAT-G12L-CS-EGFP and LAT-P17A-CS-EGFP stained with the plasma membrane (PM) and Golgi apparatus (GA) marker lectin-HPA AF647. Single-channel (EGFP and GA + PM marker) or two-channel overlay images (merge; EGFP – green, GA + PM – red, overlay – yellow) are shown together with a brightfield image (BF). Scale bars, 10 μm. More examples in Supplementary Figs. S4–S7. Representative images of 3 measurement days are shown.

### Mutation of helix-breaking amino acids cannot rescue signalling defect caused by the absence of LAT palmitoylation

Next, we analysed the effects of Pro->Ala or Gly->Leu mutations in TMD on LAT function in T cells. It was shown previously that antigen response of T cells is LAT-dependent process. Intracellular calcium mobilisation is a key part of this process and is abrogated in the absence of LAT [37]. Reconstitution of LAT-deficient J.CaM2.5 cells with transiently transfected LAT-WT resulted in high level of cytosolic calcium mobilisation after stimulation with anti-TCR antibody as measured by flow cytometry (Fig. 4A; see also **Methods**). A three-fold signal increase was observed less than a minute after stimulation, as detected by changes in fluorescence properties of the calcium-sensitive probe Fura Red (Fig. 4B). Similarly, stimulation of cells expressing LAT-P8A, LAT-G12L or LAT-P17A variants was associated with a rapid, strong, and sustained calcium response (Fig. 4B). Together, these data indicate that mutation of helix-breaking residues does not affect function of LAT in T cells.

**Figure 4.**
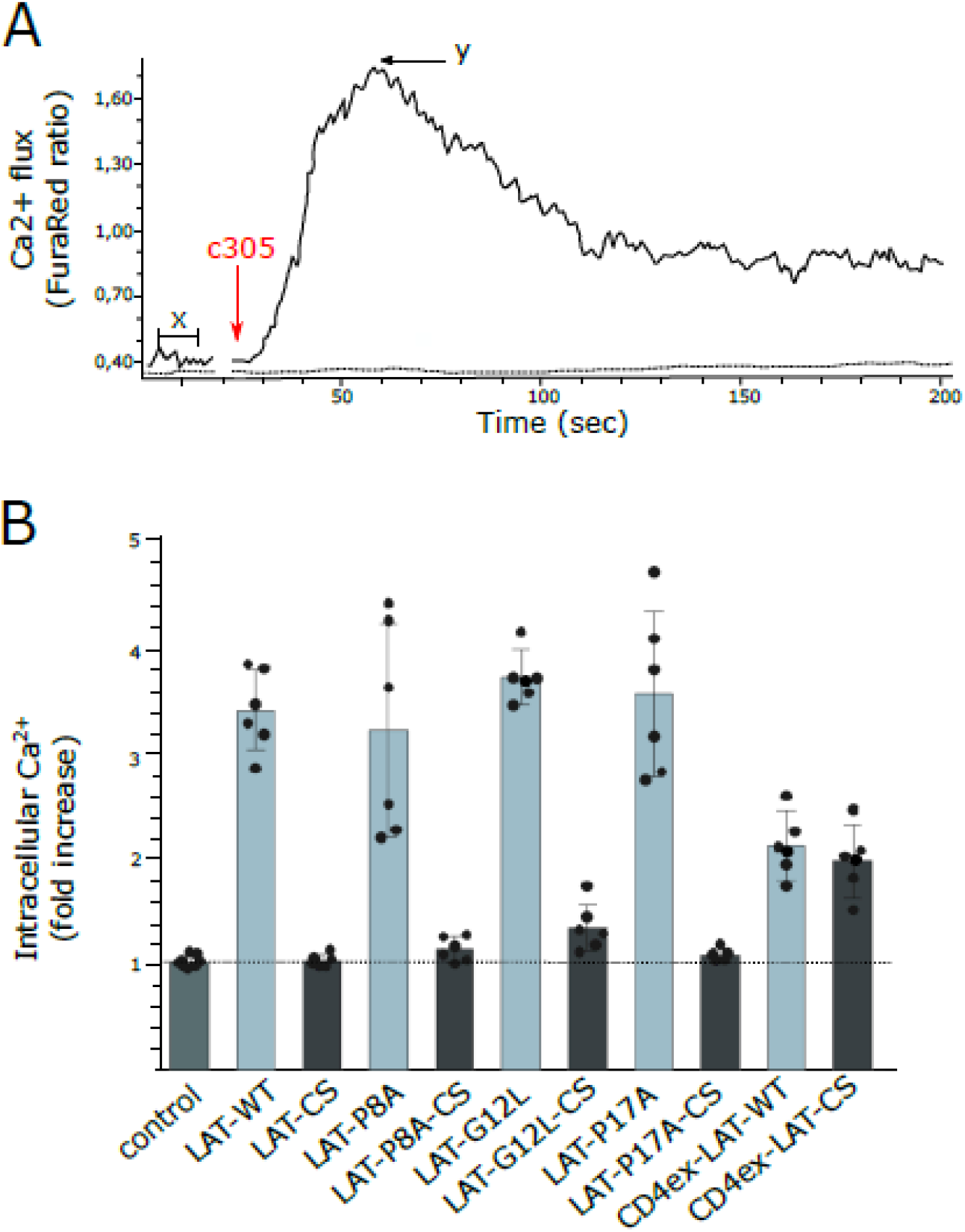
TCR-mediated calcium response of J.CaM2.5 T cells expressing LAT variants. **(A)** Representative plot of calcium mobilisation (FuraRed Ex405/Ex488) in the presence (full line) or absence (dotted line) of stimulating antibody recorded between 0 and 200 s in transiently transfected J.CaM2.5 T cells by flow cytometry. Transfected cells were stained with FuraRed and stimulated by the addition of c305 antibody 25 seconds after the beginning of the measurement (indicated by red arrow; y – maximum peak response, x - mean of the non-specific calcium flux without stimulation). For the analysis, transfected cells were selected using gating for GFP signal. **(B)** A bar graph showing increased intracellular calcium levels after TCR antibody stimulation in J.CaM2.5 T cells expressing indicated LAT variants (palmitoylatable LAT variants – dark grey bars; non-palmitoylatable LAT variants – light grey bars). Error bars indicate standard deviation of the mean, black puncta represent individual experiments. The data show measurements for 6 independent measurement days.

Interestingly, when analysing response to TCR antibody stimulation in J.CaM2.5 cells expressing any of the non-palmitoylated LAT-CS variants, no calcium mobilisation was observed regardless of the mutations in the LAT transmembrane domain (Fig. 4B). Surface expression is thus not sufficient for LAT function in antigen-induced T cell signalling. Previously, we demonstrated that addition of the CD4 extracellular domain to LAT-CS rescued plasma membrane sorting of CD4ex-LAT-CS variant ([6] and Supplementary Fig. S8). Importantly, robust calcium mobilisation is induced by antibody stimulation in J.CaM2.5 cells expressing CD4ex-LAT or its non-palmitoylatable variant - CD4ex-LAT-CS (Fig. 4B). CD4ex-LAT and CD4ex-LAT-CS contain intact LAT TMD (apart from cysteine residues 26 and 29 in the CS variant). These data indicate that replacement of helix-breaking amino acids in the TMD with helix supporting ones partially rescues LAT sorting to the plasma membrane, but such surface LAT molecules remain inaccessible for early T cell signalling molecules (e.g. Lck and ZAP-70; ref. [32]).

### Nanoscopic organisation of LAT mutants on the surface of T cells

We were thus interested whether this is due to the different nanoscopic organisation of LAT-WT and the mutants, which cannot be visualized by standard confocal microscopy. We employed super-resolution (SR) microscopy called photoactivation localization microscopy (PALM) to image nanoscopic organization of the variants LAT-WT, LAT-P17A and LAT-P17A-CS at the surface of transiently transfected J.CaM2.5 cells. For this purpose, GFP tag was replaced with mEos2, which exhibits photoconvertible behaviour essential for PALM imaging [38]. Transfected cells were immobilized on glycine-coated coverslips to avoid artificial stretching of T cells caused by poly-*L*-lysine coated surfaces [39]. As shown in Fig. 5A, LAT-WT exhibits a random distribution over the surface of unstimulated T cells. LAT-P17A variant, which generates similar calcium response in antibody-stimulated T cells, is also randomly distributed (Fig.5A, middle panel). For J.CaM2.5 cells expressing the LAT-P17A-CS variant, which do not respond to antibody stimulation, irregular distribution of molecules was observed using PALM (Fig. 5A, right panel). This is probably caused by reduced spreading of LAT-P17A-CS-transfected cells on glycine-coated coverslips as represented by a cell footprint on the optical surface analysed using interference reflection microscopy (IRM; Fig. 5B, right panel). The IRM images of these cells are highly non-homogenous indicating an extensive three-dimensional surface structure probably caused by the accumulation of microvilli or membrane ruffles at the contact site. Intensive spreading was observed for cells expressing LAT-WT and LAT-P17A (Fig. 5B). Similar protein distributions were detected for palmitoylated and non-palmitoylated LAT-P8A variants (Supplementary Fig. S9). These data indicate that mutations to Pro8 and Pro17 of the TMD do not produce alterations in the observable nanoscopic organisation of LAT at the T-cell surface. However, the ability of T cells to spread on non-stimulating surface is affected by the palmitoylation state of LAT.

**Figure 5.**
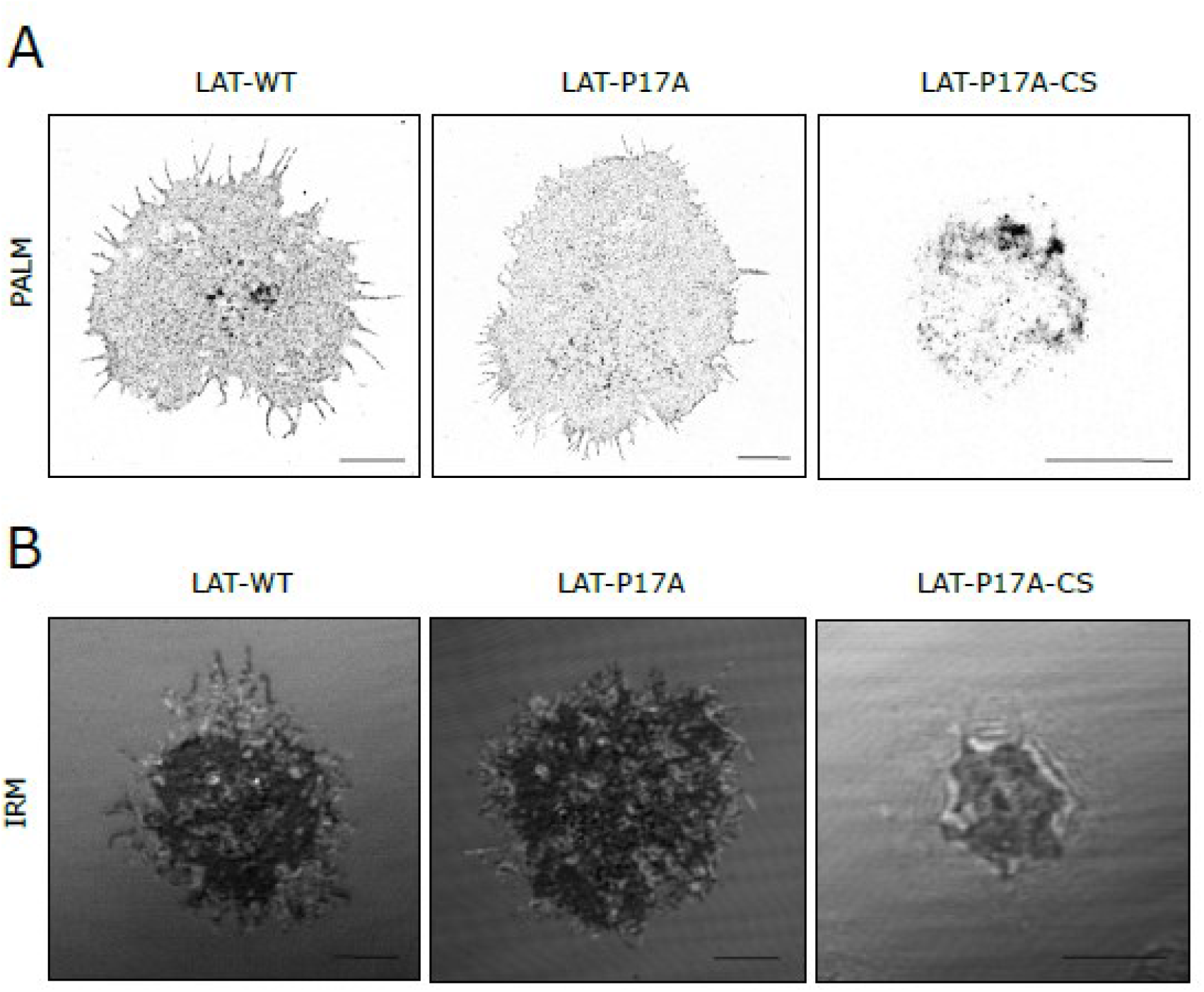
Super-resolution (SR) imaging of LAT variants on the surface of T cells. **(A)** SR images of LAT-WT, LAT-P17A and LAT-P17A-CS in transiently transfected J.CaM2.5 T cells imaged by photoactivation localisation microscopy (PALM) of mEos2 fusion proteins. The images represent a footprint of fixed cells on glycine-coated coverslips. **(B)** IRM images of living J.CaM2.5 T cells transiently transfected with LAT-WT, LAT-P17A and LAT-P17A-CS. Dark areas represent a cell contact with the optical surface. Light grey diffraction rings indicate complex surface morphology of an imaged cell. Scale bars, 5 μm. Representative images from 2 measurement days. At least 10 cells were analysed for each LAT variant.

## DISCUSSION

In this work, we demonstrate the impact of helix-breaking residues on dynamic geometry of LAT TMD and localisation of LAT in T-cell membranes. Using MD simulations, we uncovered the presence of a kink in the TMD of human LAT. The *in silico* and live cell imaging data indicate that the kink is primarily caused by the presence of proline residue in the central segment of the TMD. Replacement of this residue with alanine (P17A mutant) completely removed the kink in the LAT TMD structure. Moreover, non-palmitoylated LAT-P17A mutant localised to the plasma membrane in approximately half of the cells. No such localisation was observed for non-palmitoylated LAT with native TMD sequence, thus, indicating that Pro17 is critical for the dependence of LAT surface expression on palmitoylation. This is probably caused by a highly dynamic central region of native TMD. Indeed, similar helix destabilisation was observed in other TM peptides with centrally positioned proline residue [18]. In oligomeric and multispanning (polytopic) proteins, central prolines were found to facilitate tight packing of TMDs and to function as transmission elements of conformational changes required for a rapid response to external stimuli (e.g., ligands, photons or electrochemical fluctuations; refs. [19, 21, 22, 40]). Potentially, such kinked TMD can affect the stability of single-spanning protein such as LAT in membranes. We did not observe any cytosolic expression of native and mutant LAT protein, independent of its palmitoylation state.

TMD kinks caused by centrally positioned prolines are controlled by neighbouring amino acids [21, 22]. Our live cell imaging data, LAT localisation in T cells, indicate such role for glycine positioned five residues upstream of the central proline. However, no effect on the kink and its dynamics was observed in MD simulations. This may be caused by a subtle effect of glycine residue on TMD dynamics, which is more pronounced in the complex lipid mixture of cell membranes. Glycine was found to enhance the helix-destabilising effect of proline in channel-forming peptide alamethicin [27]. On its own, it destabilises helical structure of downstream residues, especially in the presence of additional helix-breaking residues in the TM segment [24]. The observed kink of LAT TMD spreads over 3 residues downstream of Gly12 and positions i−2 to i−4 with respect to Pro17. This is in agreement with reported destabilising effect of glycine and proline downstream and upstream, respectively, of their position in the helix [24, 35].

Interestingly, as demonstrated by MD simulations, replacement of Pro8 in the luminal part of LAT TMD for alanine increased dynamics of the kink in the central part. Moreover, non-palmitoylated LAT-P8A mutant localised to the T-cell surface in a similar proportion of cells as observed for the LAT-P17A mutant. A minor reduction of the helicity of native LAT TMD was observed around Pro8 residue. However, similar instability was detected for all tested peptides, including P8A variant. This minor helical instability is probably not caused solely by Pro8. Von Heijne and Deber groups reported significant impact of prolines localised at the TMD ends on protein (peptide) function and structure [41, 42]. On the contrary, no effect of central proline on TM helix was found using protein topology-sensitive glycosylation assay [41]. We have observed a strong impact of central proline and only a minor, if any, of proline present in the first loop of the helix. We speculate that local lipid environment and overall structure of the TM segment modulate impact of TM prolines on membrane proteins.

Cellular membranes can have diverse lipid composition, which, among other things, determines their thickness [43]. Hydrophobic segment of human LAT TMD is rather long (24 residues, or 23 residues as reported in uniprot.org) but the protein requires palmitoylation for its sorting to the thicker bilayer of the plasma membrane [6, 33, 34]. This contradicts previous observations that, in vertebrates, TMDs with >22 residues are sufficient for protein sorting to the plasma membrane [8, 9]. It is therefore possible that the kink modulates the apparent ‘length’ of LAT TMD by adapting the helix fold to the local lipid environment.

Recently, Levental and colleagues reported the role of TMD asymmetry in protein sorting to the plasma membrane [44]. Using bioinformatic approach, they found that proteins at the plasma membrane have smaller surface area in the luminal (exoplasmic) half of their TMD compared to the cytosolic half. This is in agreement with previously reported accumulation of voluminous amino acids in the cytosolic part of the TMD of the plasma membrane proteins [9, 10]. The authors argue that chemical asymmetry of the plasma membrane, which exhibits higher rigidity in its outer leaflet compared to the inner leaflet, selects proteins with asymmetric TMD in terms of surface area [44]. Voluminous amino acids are randomly distributed over the sequence of LAT TMD. LAT TM segment thus does not exhibit asymmetry with respect to the surface area. We have previously shown that voluminous, non-aromatic amino acids form a rough surface of TM segments and can locally rigidify membranes [45]. Strong presence of voluminous, non-aromatic amino acids in LAT TMD (14 out of 24 residues) and their distribution may, thus, require increased flexibility of this segment to avoid interference with the plasma membrane function, which exhibits lipid composition prone to rigidify.

Indeed, LAT is one of very few integral membrane proteins, which segregate into more ordered parts of membranes, at least in giant plasma membrane vesicles [46]. Even though, membrane domains with similar properties have not been indisputably proven in cells yet, their existence is supported by extensive indirect evidence [43, 47]. The ability of LAT TMD to rapidly change geometry (intrahelically) may represent an advantage for its inclusion into membrane areas with more rigid organisation. Stabilisation of such structures is predicted to involve curvature [48]. The kink in LAT TMD can again help to localise the protein into membranes with complex geometry.

In summary, we demonstrate the existence of kink in LAT TMD, which controls LAT sorting to the plasma membrane. The kink is induced by centrally positioned proline but is further modulated by neighbouring glycine and proline. Replacement of these residues with helix-stabilising amino acids partially recovers LAT sorting to the plasma membrane but not the function of this signalling adapter. These data indicate a complex control of LAT function by highly dynamic structure of its TMD.

## ACKNOWLEDGEMENT

The work was supported by Charles University Grant Agency (GAUK; project number 298216) and Czech Science Foundation (19-26854X). It also received institutional funding from IMG CAS, Prague, Czech Republic (RVO 68378050). We acknowledge Light Microscopy Core Facility, IMG CAS supported by MEYS (LM2015062, CZ.02.1.01/0.0/0.0/16_013/0001775), OPPK (CZ.2.16/3.1.00/21547) and MEYS (LO1419).

## METHODS

### Cell culture and transfection

LAT-negative Jurkat T cell variant (J.CaM2.5) [49], kindly provided by Art Weiss (University of California San Francisco), was cultured in RPMI-1640 media (Sigma-Aldrich), complemented with 10% fetal calf serum (FCS; Life Technologies) in a humidified incubator (Eppendorf) under controlled conditions of 37°C and 5% CO_2_. The cells were transiently transfected using the Neon^®^ transfection system (Life Technologies) according to the manufacturer's instructions. One μg of vector DNA was used per shot (3 pulses of 1325 V for 10 ms) per 500 000 cells, which were then incubated in 0.5 ml of media. Cells were analysed 16-20 hours post transfection.

### DNA cloning

To generate LAT P8A and G12L mutants, LAT TMD coding sequence was modified using polymerase chain reaction (PCR) and the primers with the appropriate mutations (Supplementary Table S1), and cloned into pXJ41-LAT-EGFP plasmid (or its non-palmitoylatable CS mutant) using EcoRI and BamHI restriction sites. The variants without the CD148 leader, c-Myc and 5′ UTR were used [6]. LAT-P17A variants were prepared by annealing the appropriate oligonucleotides (Supplementary Table S1) and subcloning as described above. For super-resolution microscopy, the EGFP coding sequence was changed to mEos2 using BamHI and XhoI restriction sites.

### Live-cell confocal microscopy

Cells transiently transfected with GFP variants of LAT in pre-heated, colour-free RPMI-1640 medium supplemented with 10% FCS were placed on 1% gelatine-coated Ibidi μ-Slide 8-well chambers (Ibidi, Germany) and imaged using a Leica TCS SP8 laser scanning confocal microscope equipped with sensitive hybrid detectors (HyD), 488 nm (20mW) and 638 nm (30 mW) lasers and a 63×1.4 NA oil-immersion objective. Five to ten percent laser power and 10% gain were used for image acquisition. Acquired images were manually thresholded to remove signal noise detected outside of the cell using ImageJ software package [50].

### Two-colour confocal microscopy of fixed cells

Cells were immobilized on poly-*L*-lysine-coated Ibidi μ-Slide 8-well chambers for 5 minutes, fixed with 4% paraformaldehyde in PBS for 20 minutes at room temperature and subsequently washed with PBS. Cells were permeabilized for 40min using 10x Permeabilisation Buffer (eBioscience) diluted in water, washed with PBS, and subsequently blocked with 5% BSA. Plasma membrane and Golgi apparatus of transfected cells were stained with 0.5μg/ml lectin-HPA conjugated to Alexa Fluor 647 in PBS supplemented with 1% BSA (Life Technologies) and washed twice with PBS prior imaging. Images were taken using a Leica TCS SP8 laser scanning confocal microscope equipped with a 63× 1.4 NA oil-immersion objective. Minor contrast and/or brightness level adjustments were applied, and images were processed for publishing by ImageJ software.

### Single Molecule Localization Microscopy (SMLM)

#### Sample preparation

Round 25 mm coverslips (No 1.5H, High precision; Marienfeld) cleaned with Helmanex III solution, were coated with 500 μl of 2M glycine solution in ultrapure water for 30 minutes at room temperature. Prior to landing of the cells, coated coverslips were washed with Mili Q^®^ water once (Merck Millipore). Transfected cells were harvested (3 min at 300 rcf), transferred into PBS pre-warmed to 37°C and added to the coated coverslips. Cell immobilization was achieved by incubation at 37°C for 10 minutes. Afterwards, cells were fixed for 1 hour at room temperature with 4% paraformaldehyde (Electron Microscopy Sciences) in PBS containing 2% sucrose (Sigma-Aldrich). Fixation was stopped with 50mM NH_4_Cl in PBS followed by three rounds of washing with PBS. Coverslips with cells were transferred into ChamLide holder (LiveCell Instruments). 200 nm gold beads (fiducial markers; BBI) were loaded to the sample in 0.9% NaCl for 5 minutes. Imaging was performed in PBS.

#### Microscope setup and measurements

SMLM measurements were performed on a home built microscope (IX71 body; Olympus) equipped with 150mW 405nm (Cube; Coherent) and 150mW 561nm (Sapphire; Coherent) lasers, 100× 1.49 NA objective (UApoN; Olympus), EMCCD camera (iXon DU-897, Andor) and manual TIRF tilting mechanism (Thorlabs). Synchronisation of laser switching, and camera recording was performed with two acousto-optic tuneable filters (AOTF; AOTFnC-400.650-TN, AA Optoelectronics) and a home written acquisition software (LabView). To turn the fluorophores into the dark state, fast epifluorescence irradiation with 561nm laser was applied. Data were acquired using the highly inclined and laminated optical sheet (HILO) illumination mode with 50 ms/frame collection time. The power of activation laser (405 nm) was increased gradually to collect 10 000 – 15 000 frames with a uniform emitter signal.

#### Data analysis

To generate super-resolution maps of LAT localisations, data were analysed using ThunderSTORM plugin of the ImageJ/Fiji software package [50].

### Interference Reflection Microscopy (IRM)

IRM of living cells immobilized on glycine-coated coverslips was recorded on a modified Olympus Fluoview 1000 setup equipped with 60× 1.2 NA water immersion objective (UPlanSApo; Olympus). GFP signal was excited with 20mW 488nm laser (Sapphire; Coherent) and recorded on a single-channel PMT detector equipped with a selective dichroic mirror (DM488/543/633; Olympus). For acquisition and system control the FluoView 1000 software package (Olympus) was used. The images were processed using ImageJ/Fiji software package [50].

### Calcium measurements by flow cytometry

Cells resuspended at 10^6^ cells/ml in pre-warmed PBS containing 1 μM Fura Red AM (Invitrogen) were incubated at 37°C under 5% CO_2_ in a humidified incubator for 30 minutes. Cells were then washed twice with PBS, resuspended in RPMI-1640 (Sigma-Aldrich) supplemented with 10% FCS (Life Technologies) and kept on ice. Prior to measurement, cells were equilibrated to 37°C for 4 minutes in a water bath.

All measurements were performed on an LSRII flow cytometer (BD Bioscience). A steady-state (background) signal was recorded for 25 seconds. Afterwards, the cells were stimulated with an antibody: pre-warmed RPMI (Sigma-Aldrich) supplemented with 10% foetal calf serum (Life Technologies) containing anti-TCR antibody C305 (home-produced supernatant, 10 μg/ml) was added in 1:1 ratio. Changes in calcium level were recorded continuously for over 200 seconds. Transfected cells were gated for EGFP positivity. The relative calcium concentration was measured as a ratio of the Fura Red fluorescence intensity elicited by the excitation at 405 nm (emission measured at 635–720 nm) and 488 nm (emission measured at 655–695 nm). The calcium values were calculated as the increasing signal stimulated by the 405 nm laser over the decreasing signal stimulated by the 488 nm laser using the Kinetics tool in FlowJo software version 10.6.1 (Tree Star Inc., OR, USA). The fold increase in signal was calculated as a ratio between maximum peak response and a mean of the steady-state signal recorded without stimulation, as indicated in Fig. 4.

### All-atom MD simulations

For atomistic MD simulations, the amino acid sequence of wild type LAT (EEAILVPCVLGLLLLPILAMLMALCVHCHRLP) was used to construct a fully helical model of the peptide employing the Protein Builder of the Molefacture plugin in the VMD software [51]. The standard GROMACS tool (pdb2gmx) was used to apply the fully atomistic AMBER99SB-ILDN force field for the peptide [52, 53]. A lipid membrane consisting of 128 POPC molecules (64 in each leaflet) was hydrated with ~5000 water molecules and pre-equilibrated. The Slipid force field was used for phospholipids and TIP3P model was employed for water [54–56]. Then, the method of Javanainen was employed to insert the LAT peptide into the membrane in a transmembrane orientation [57]. The overall −1 charge of LAT was neutralized by inserting one Na^+^ cation into the water phase. The standard Dang’s force field parameters were used for the sodium cation [58]. The membrane with LAT was then simulated using GROMACS software [52]. The same protocol of system preparation and simulations was used for the three LAT mutants considered in MD.

In simulations, fully periodic boundary conditions were used. The simulations were performed at the temperature of 310 K controlled by Nose-Hoover thermostat with the time constant 1.0 ps [59]. Pressure of 1 bar was controlled employing the semi-isotropic Parrinello-Rahman barostat with the time constant of 5 ps and compressibility of 4.5×10^−5^ bar^−1^ (ref. [60]). The Particle Mesh Ewald algorithm was used for accounting for long-range electrostatic interactions with the real-space cut-off set to 1.2 nm [61]. For long-range non-bonded interactions, 1.2 nm cut-off was used, switched at the distance of 1.0 nm. All covalent bonds were constrained by LINCS algorithm, and water molecules were constrained by SETTLE method [62, 63]. The equations of motion were integrated with 2 fs time step. All simulations were performed for 1000 ns. Stabilization of the root mean square displacement and peptide-lipid contacts was used as an equilibration criterion. The initial 500 ns were treated as equilibration period while the final 500 ns were used for the analysis.

The analysis of the simulated trajectories was performed using standard GROMACS tools combined with in house Python scripts. In the case of the helix tilt angle and bending, the method of Bansal et al. implemented in the HELANAL Python library was applied [64]. The numerical data were presented using Matplotlib Python library and MATLAB [65]. Molecular visualization was performed with the VMD software [51].

## Supplementary Information

**For the manuscript:** Glatzova et al. The role of prolines and glycine in the transmembrane domain of LAT

**Supplementary Figure S1.**
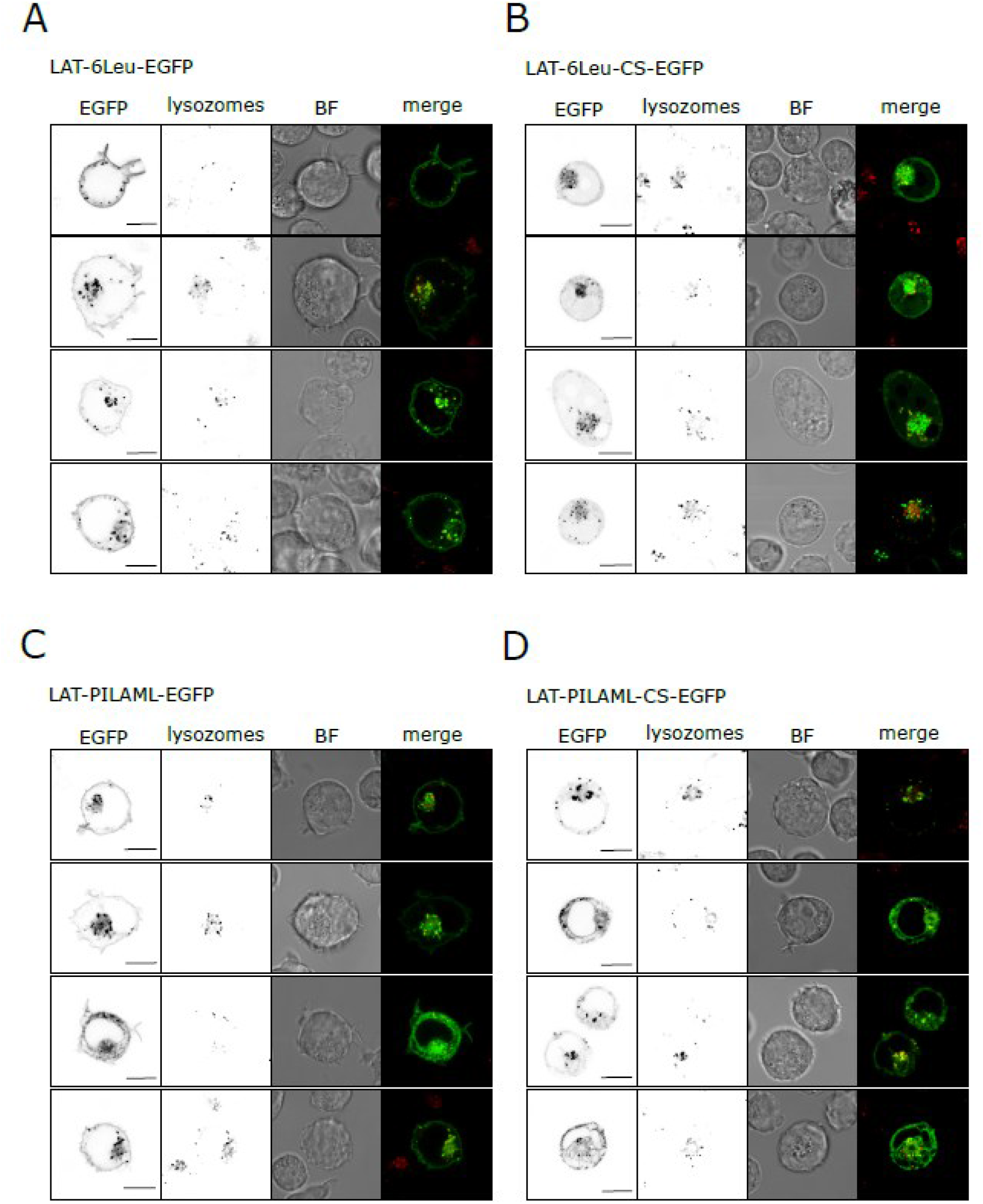
Localisation of LAT variants with prolonged TMD in J.CaM2.5 T cell line. Confocal microscopy of J.CaM2.5 T cells expressing LAT-6Leu-EGFP **(A)**, LAT-6Leu-CS-EGFP **(B)**, LAT-PILAML-EGFP **(C)** and LAT-PILAML-CS-EGFP **(D)** stained with the marker of lysosomes (Lysotracker-Red). Single-channel (EGFP and Lysosome marker) or two-channel overlay images (Merge; EGFP – green, Lysosome – red, overlay – yellow) are shown together with a brightfield image (BF). Scale bars, 10 μm.

**Supplementary Figure S2.**
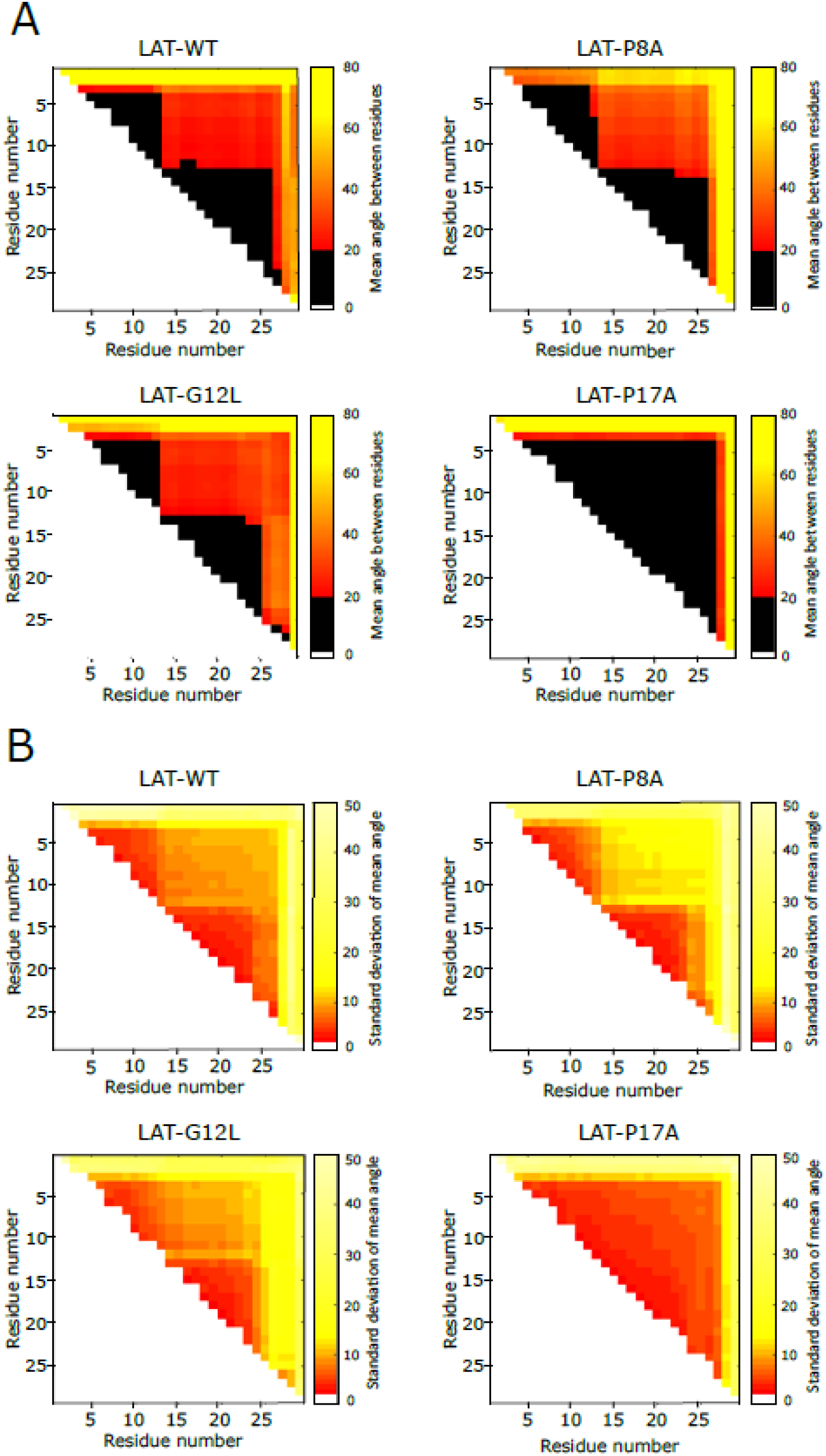
Angles formed by kinked helix of LAT peptides in POPC membrane – MD simulation data. **(A)** Plots representing mean angle between individual amino acids (Residues) of the LAT WT peptide and indicated mutant variants. Angles were calculated by HELANAL Python library with the geometry of the α-helix characterized by computing local helix axes and local helix origins for four contiguous carbon-α atoms. Angles larger than 20° indicate non-helical structure [64]. **(B)** Plots representing standard deviation of mean angles between individual amino acids (as in A) of the LAT WT peptide and indicated mutant variants. Higher values of standard deviation indicate larger flexibility of the TMD peptide within kinked region.

**Supplementary Figure S3.**
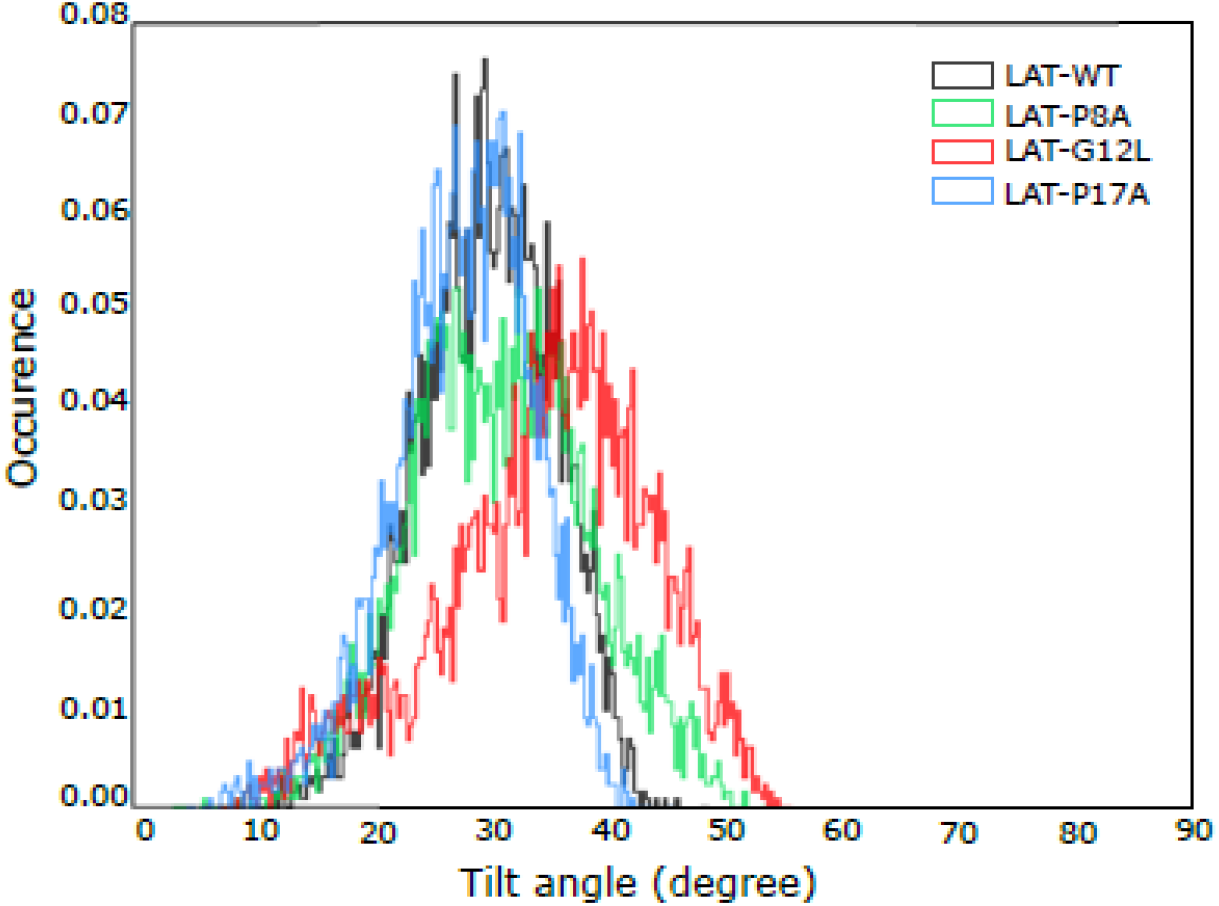
Tilt angles of LAT peptides in POPC membrane – MD simulation data. **(A)** A plot of tilt angles with respect to the membrane normal accommodated by the LAT WT peptide and indicated mutant variants. The data represent a distribution of tilt angles from the trajectories of the peptides in POPC membranes acquired in all-atom MD simulations. Tilt angles were calculated by HELANAL Python library [64]. The tilt angle value of zero represents a helix orientation parallel to membrane normal, and the tilt angle of 90 degree – perpendicular to the membrane normal.

**Supplementary Figure S4.**
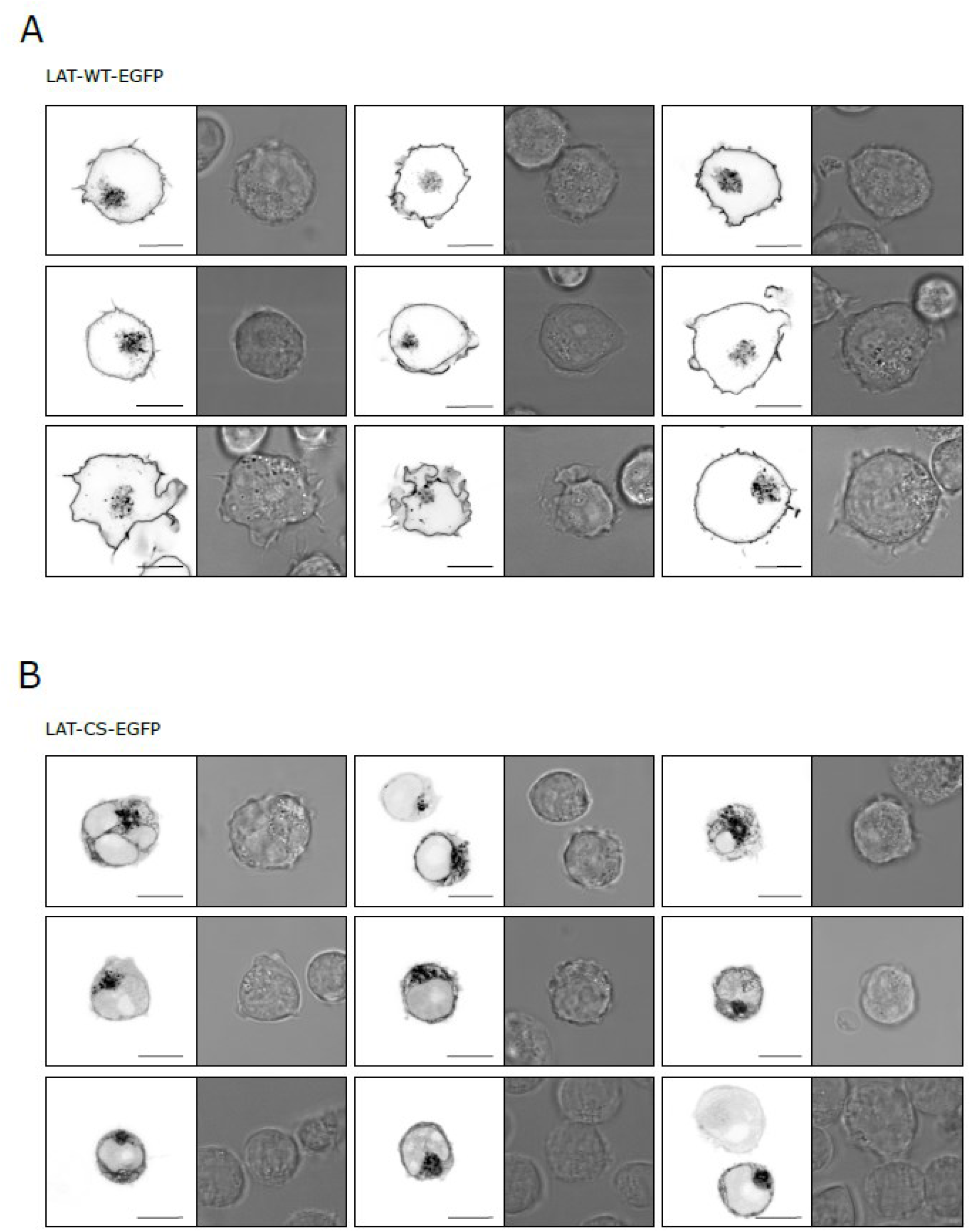
Localisation of LAT-WT-EGFP (A) and its non-palmitoylatable variant LAT-CS-EGFP (B). Nine examples of J.CaM2.5 T cells expressing LAT variants imaged by confocal microscopy. EGFP channel is shown along the brightfield image. In total, 74 cells were analysed for LAT-WT and 51 cells for LAT-WT-CS variant. Scale bars, 10 μm.

**Supplementary Figure S5.**
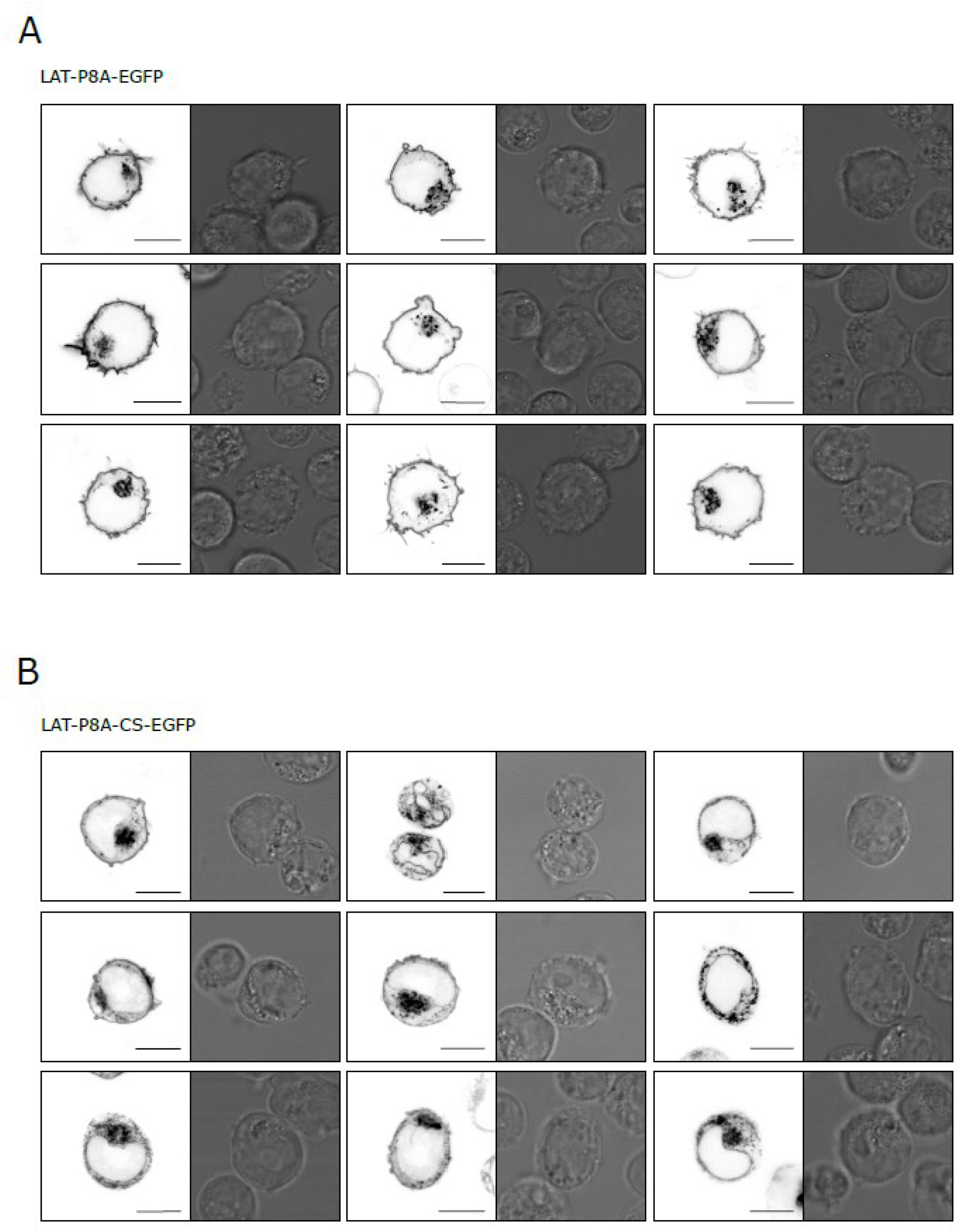
Localisation of LAT-P8A-EGFP (A) and its non-palmitoylatable variant LAT-P8A-CS-EGFP (B). Nine examples of J.CaM2.5 T cells expressing LAT variants imaged by confocal microscopy. EGFP channel is shown along the brightfield image. In total, 80 cells were analysed for LAT-P8A and 76 cells for LAT-P8A-CS variant. Scale bars, 10 μm.

**Supplementary Figure S6.**
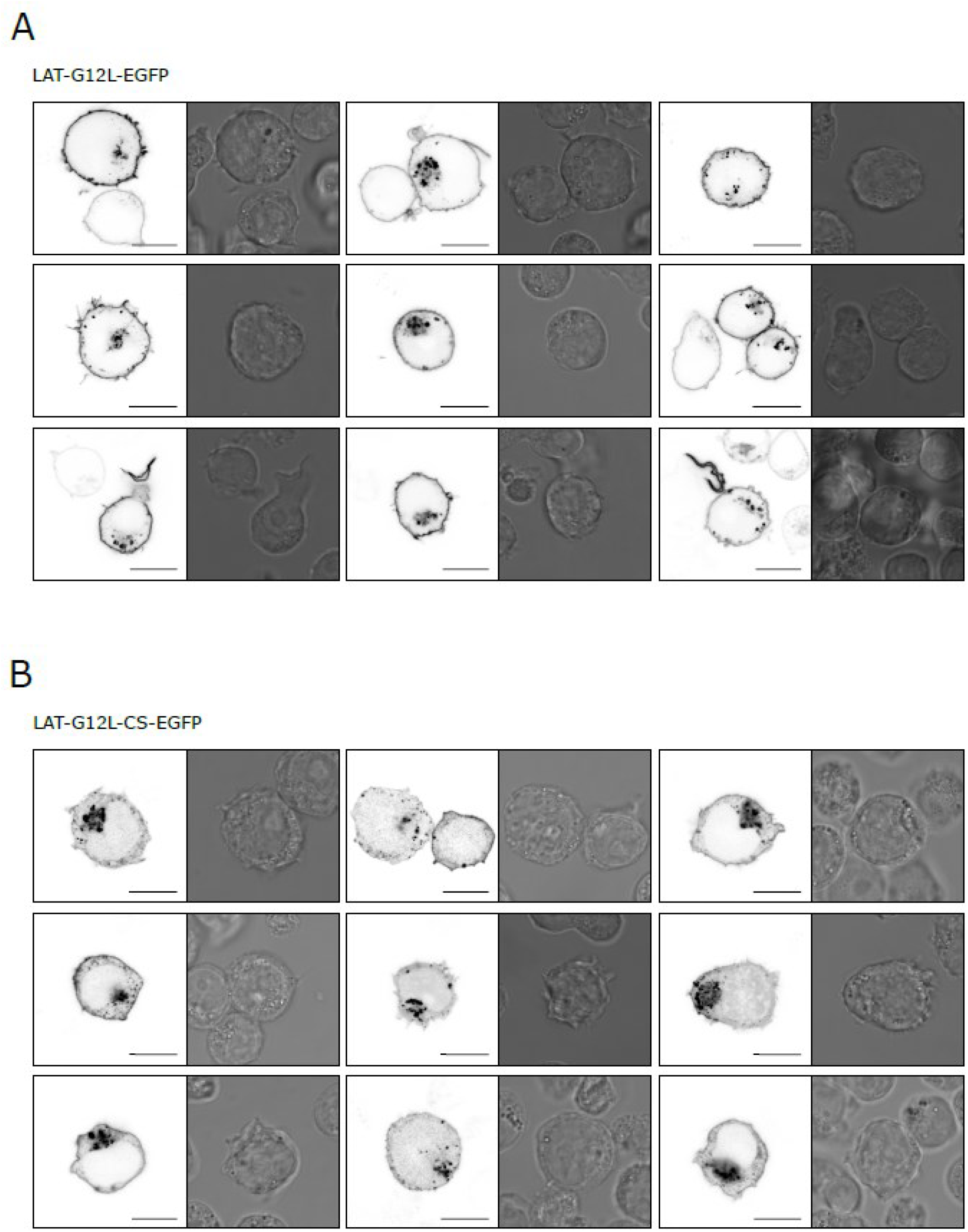
Localisation of LAT-G12L-EGFP (A) and its non-palmitoyable variant LAT-G12L-CS-EGFP (B). Nine examples of J.CaM2.5 T cells expressing LAT variants imaged by confocal microscopy. EGFP channel is shown along the brightfield image. In total, 90 cells were analysed for LAT-G12L and 55 cells for LAT-G12L-CS variant. Scale bars, 10 μm.

**Supplementary Figure S7.**
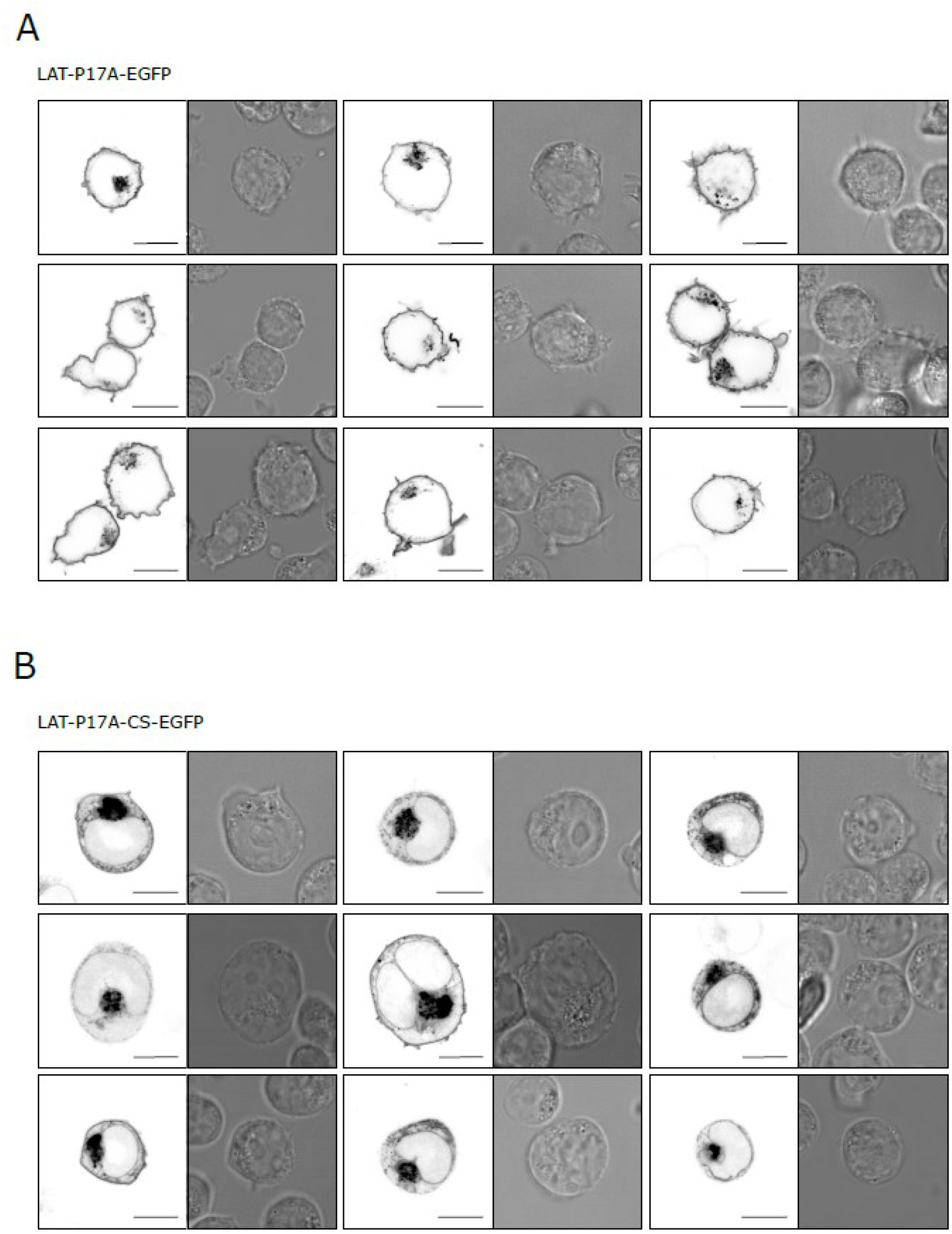
Localisation of LAT-P17A-EGFP (A) and its non-palmitoylatable variant LAT-P17A-CS-EGFP (B). Nine examples of J.CaM2.5 T cells expressing LAT variants imaged by confocal microscopy. EGFP channel is shown along the brightfield image. In total, 118 cells were analysed for LAT-P17A and 56 cells for LAT-P17A-CS variant. Scale bars, 10 μm.

**Supplementary Figure S8.**
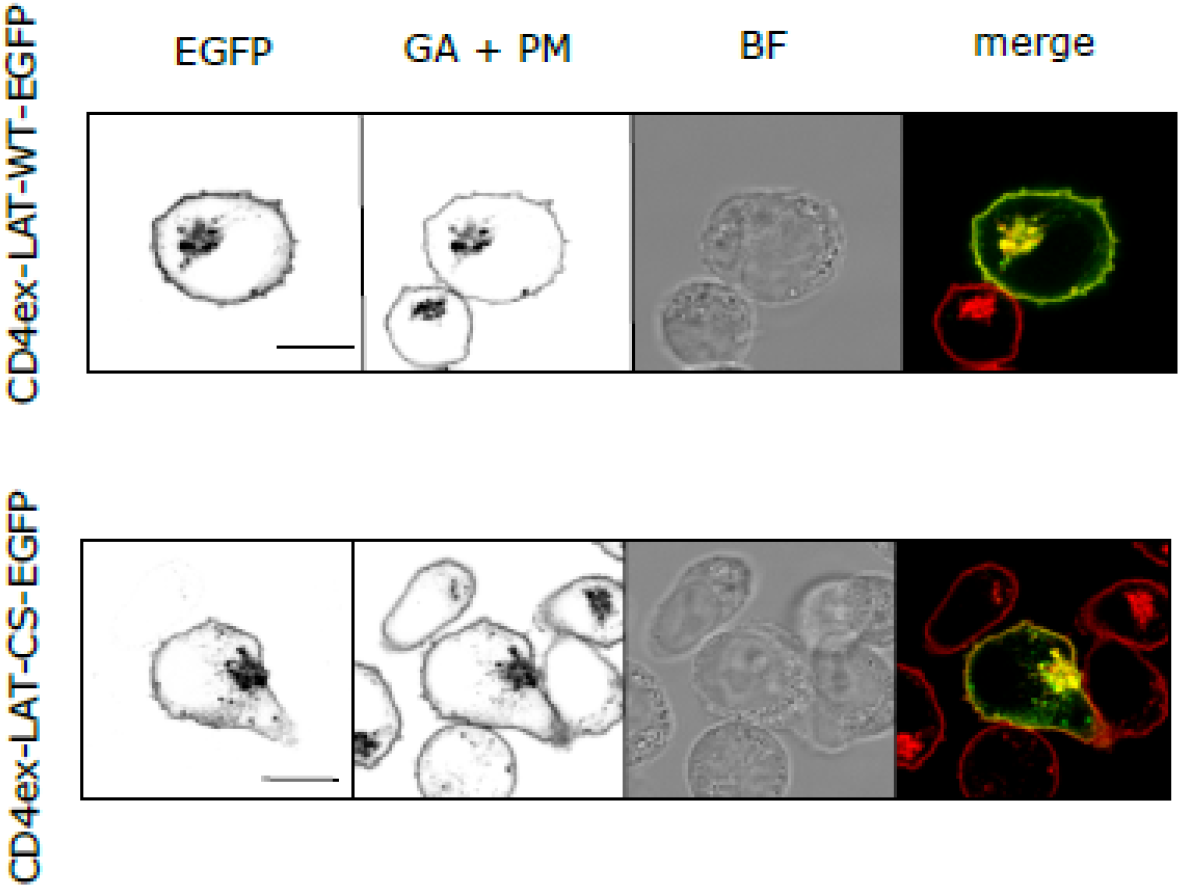
Localisation of LAT variants containing the extracellular domain of CD4 in J.CaM2.5 T cell line. Confocal microscopy of J.CaM2.5 T cells expressing CD4ex-LAT-EGFP **(upper panel)** and CD4ex-LAT-CS-EGFP **(lower panel)** stained with the plasma membrane (PM) and Golgi apparatus (GA) marker lectin-HPA AF647. Single-channel (EGFP and GA + PM marker) or two-channel overlay images (Merge; EGFP – green, GA + PM – red, overlay – yellow) are shown together with brightfield image (BF). Scale bars, 10 μm.

**Supplementary Figure S9.**
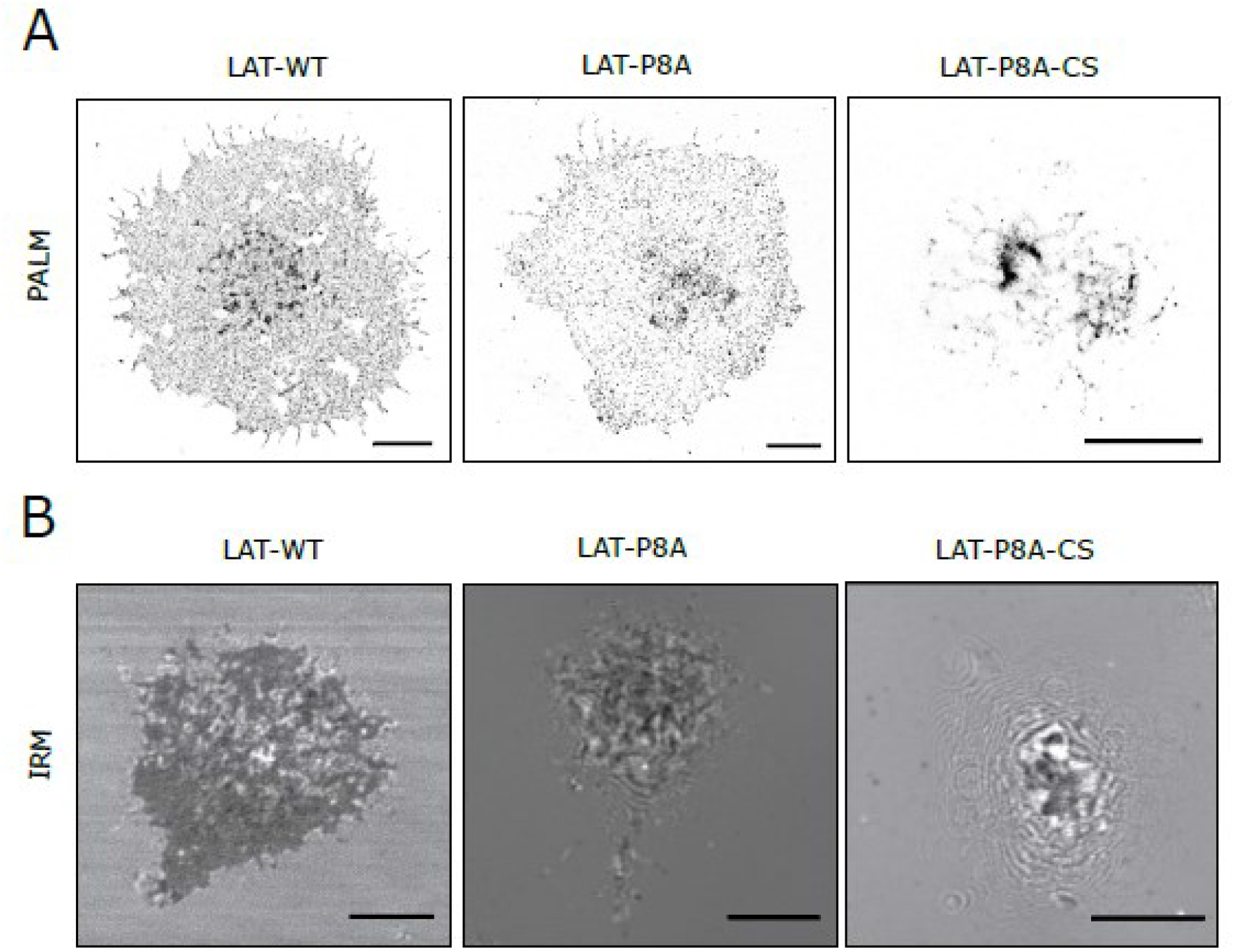
Super-resolution (SR) imaging of LAT variants on the surface of T cells. **(A)** SR images of LAT-WT, LAT-P8A and LAT-P8A-CS in transiently transfected J.CaM2.5 T cells imaged by photoactivation localisation microscopy (PALM) of mEos2 fusion proteins. The images represent a footprint of fixed cells on glycine-coated coverslips. **(B)** IRM images of living J.CaM2.5 T cells transiently transfected with LAT-WT, LAT-P8A and LAT-P8A-CS. Dark areas represent a cell contact with the optical surface. Light grey diffraction rings indicate complex surface morphology of an imaged cell. Scale bars, 5 μm. Representative images from 2 measurement days. At least 9 cells were analysed for each LAT variant.

**Supplementary Table S1.**
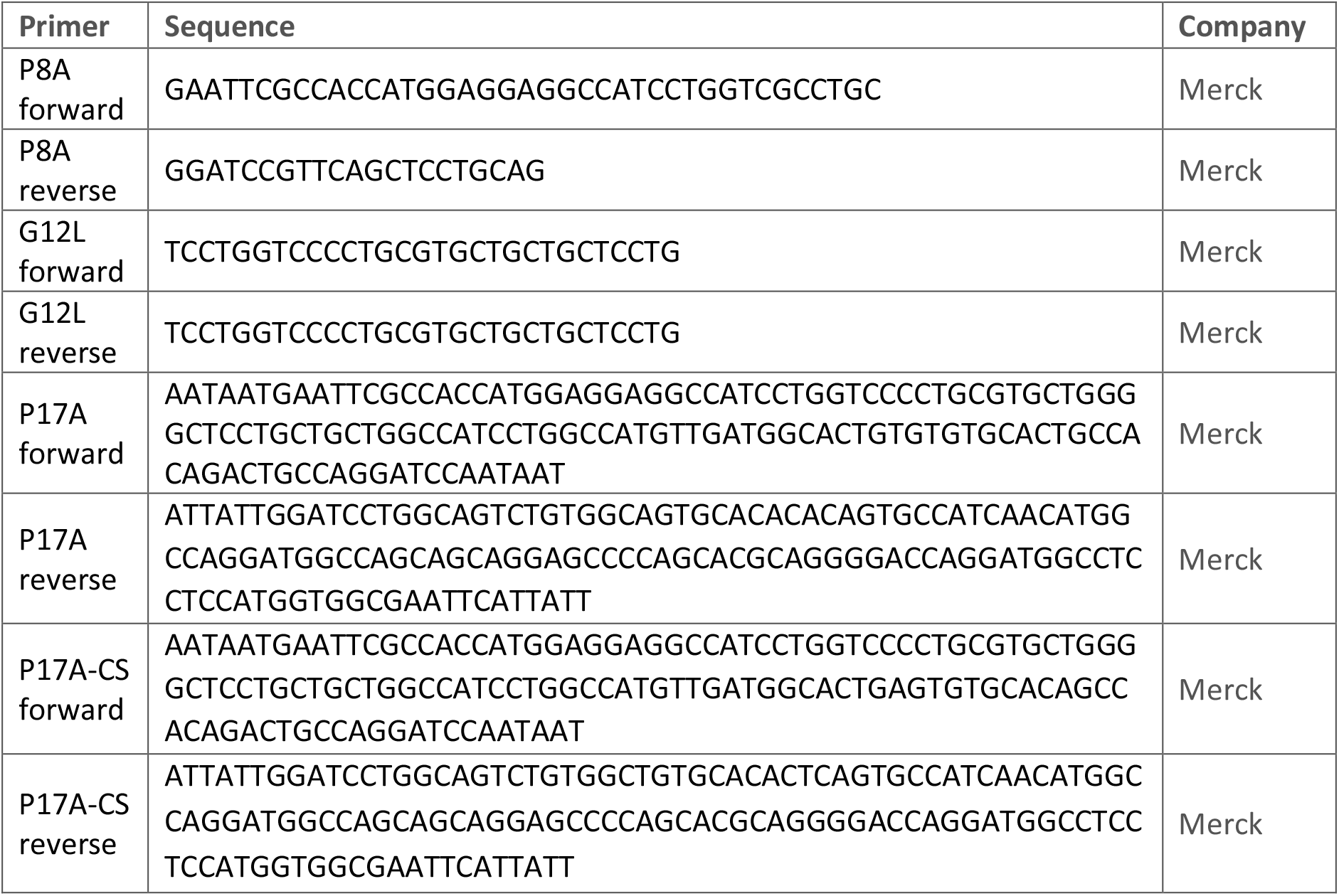
List of PCR primers.

## SUPPLEMENTARY MOVIE LEGENDS

**Supplementary Movie 1. The last 500 ns of the 1 μs-long MD simulation of the LAT WT peptide in POPC bilayer**, related to Figure 1b. Side view (parallel to the bilayer midplane) of the simulation box is shown. The video was created based on the trajectory sampled each 500 ps. The trajectory was centered with respect to the peptide center of geometry. Phosphorous atoms of POPC are shown as cyan balls, and the peptide is depicted using NewCartoon representation (purple). The remaining atoms and molecules, including water, are not shown for clarity. The video was prepared employing MovieMaker plugin of VMD. H264-MPEG-4 encoding was used.

**Supplementary Movie 2. The last 500 ns of the 1 μs-long MD simulation of the LAT P8A peptide in POPC bilayer**, related to Figure 1b. Side view (parallel to the bilayer midplane) of the simulation box is shown. The video was created as described for Movie 1.

**Supplementary Movie 3. The last 500 ns of the 1 μs-long MD simulation of the LAT G12L peptide in POPC bilayer**, related to Figure 1b. Side view (parallel to the bilayer midplane) of the simulation box is shown. The video was created as described for Movie 1.

**Supplementary Movie 4. The last 500 ns of the 1 μs-long MD simulation of the LAT P17A peptide in POPC bilayer**, related to Figure 1b. Side view (parallel to the bilayer midplane) of the simulation box is shown. The video was created as described for Movie 1.

